# AMBER: mapping accessible chromatin associations through barcode seed expansion

**DOI:** 10.64898/2026.11.18.767643

**Authors:** Robert Tan, Jinghui Song, Chenxu Zhu

## Abstract

Accessible chromatin establishes higher-order regulatory structures spanning multiple spatial scales. We developed AMBER, a hydrogel-assisted approach for confined expansion of barcode seeds and subsequent recording of accessible chromatin associations. In mouse brain cells, we characterized the multi-scale regulatory organizations that accessibility alone cannot explain. We identified a subset of genomic regions enriched for inter-chromosomal interactions that also encode proteins engaged in interaction networks. These regulatory hubs are essential for neural function and help predict disease-risk loci and regulators.

## Main

Biological organization spans a broad range of spatial scales, from nanometer-scale molecular complexes and enhancer-promoter interactions to micrometer-scale nuclear compartments and transcripts produced by individual cells, and the millimeter-scale tissue niches^1-3^. A powerful strategy to characterize the molecular constituents of such associations is to assign shared molecular identifiers to biologically related molecules. For example, pre-manufactured barcoded beads in single-cell RNA sequencing^4, 5^ and spatially patterned barcode probes^6, 7^ in spatial transcriptomics enable the identification of transcripts originating from individual cells or defined tissue locations, respectively. Alternatively, DNA microscopy exploits the diversity of randomized DNA seeds and proximity molecular interactions to reconstruct the spatial organization of molecules from their connectivity networks^8^. These approaches demonstrate the value of recording biological associations with shared molecular identifiers; however, generating multiple copies of a unique barcode while keeping them locally confined remains a key design challenge for molecular co-registration.

Accessible chromatin provides binding sites for transcriptional regulators and cofactors, which collectively establish higher-order regulatory structures spanning multiple spatial scales^9^. These structures have traditionally been investigated using complementary technologies, each capturing different aspects and scales of chromatin organization^10^. Here we introduce AMBER (<u>Am</u>plification of random barcodes for co-Embedded chromatin registration), a hydrogel-assisted strategy for confined expansion of barcode seeds and subsequent recording of molecular associations (**Fig. 1a**). AMBER’s capability for multi-molecular co-registration was validated using model DNA oligo reporters and chromatin fragments from different species, and its utility in characterizing higher-order chromatin associations was demonstrated by applying it to mouse brain nuclei.

**Fig. 1:**
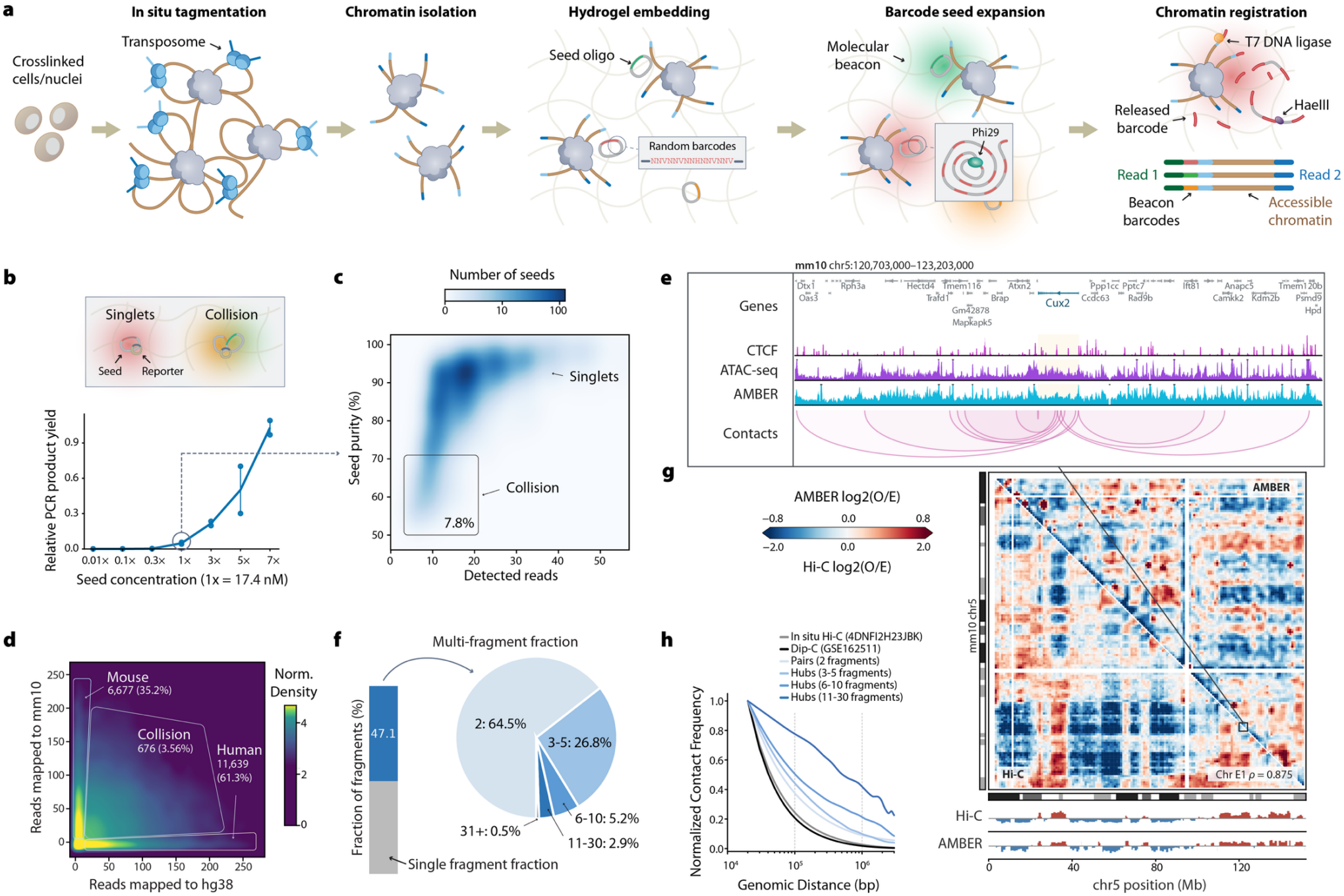
AMBER profiles accessible chromatin associations. **a**, Schematic of AMBER for mapping co-localized accessible chromatin. **b**, Quantification of beacon barcode collision in hydrogels with the dual-oligo barcoding assay (n = 2 technical replicates). **c**, Density scatter plot showing the observed seed-reporter barcode collision from sequencing. **d**, Density scatter plot showing the observed chromatin complex barcode collision from the species mixing experiment. **e**, Genome browser view of the representative AMBER signals at the Cux2 locus (mm10 chr5: 120,703,000-123,203,000). Reference tracks: ATAC-seq (ENCSR889WQX) and CTCF (ENCSR644VYX) are shown for comparison. **f**, Fraction of reads classified as singlets, pairs, and multi-way contact hubs of different sizes. **g**, Heatmap showing the contact maps of chr5 at 1Mb resolution for AMBER and Hi-C (4DNFI2H23JBK). **h**, Contact frequency decay curves relative to linear genomic distance for AMBER hubs stratified by hub size, reference in situ Hi-C (grey; 4DNFI2H23JBK) and Dip-C (black; GSE162511) are also shown.

Polyethylene glycol (PEG) and polyacrylamide hydrogels permit a wide range of nucleic acid-modifying enzymes^11^ while restricting diffusion of large oligonucleotide fragments^12^. AMBER builds on this property; the procedure begins with Tn5 tagmentation of light formaldehyde-crosslinked cells or nuclei to attach a biotinylated DNA adaptor to accessible chromatin regions. Next, nuclei are lysed, the fragmented chromatin complexes are purified, and they are co-embedded with barcode seeds into a thiol-acrylate PEG hydrogel. The barcode seed is designed as a circularized single-stranded DNA with a 15-bp random sequence flanked by restriction enzyme cleavage sites and a hybridization handle (**Extended Data Fig. 1a** and **Supplementary Table 1**). Hydrogel-confined rolling circle amplification (RCA) of the barcode seed is performed by Phi29 DNA polymerase, followed by endonuclease cleavage to release barcode monomers, generating the molecular “beacons” that can ligate to the Tn5 transposition-introduced handle on the associated chromatin fragments (**Extended Data Fig. 1c**, and **Extended Data Fig. 2a**). We found that a minimum length of 8-bp overhang sequence is required for efficient in-gel ligation and the T7 DNA ligase outcompetes T4 DNA ligase and HiFi Taq DNA Ligase (ligation efficiency: 78.01% vs. 1.54% and 23.15%, respectively). Interestingly, T4 DNA ligase generated spurious barcode concatemerizations, likely due to its ssDNA ligase activity when barcode monomers were present in molar excess. Finally, the hydrogel is dissolved by alkaline hydrolysis of acrylate ester linkers; barcoded chromatin fragments are reverse-crosslinked, purified, and amplified to give sequencing libraries.

An optimal seed concentration should balance molecular recovery (volumes covered by molecular beacons, and thus, registrable) against barcode collisions (volumes co-occupied by multiple molecular beacons). To identify the optimal seed concentration, we performed a titration using dual-oligo barcoding experiments: both DNA oligo types (seed and reporter) contain randomized sequences and are designed to hybridize and barcode each other. We found that 17.4 nM is the minimum seed concentration for effective amplification, and increasing oligo concentrations showed a proportional exponential increase in the amplification product yield (**Fig. 1b**). Beacon overlap was estimated at 7.8% based on the fraction of reporters paired with more than one seed with substantial reads detected (**Fig. 1c**). The collision rate was further confirmed by co-registration of pre-mixed human and mouse chromatin complexes (**Fig. 1d**). Importantly, chromatin co-registration throughput increased with gel size, showing that enzyme diffusion was not physically limited by hydrogel thickness (**Extended Data Fig. 2b**). Thus, AMBER is a proximity ligation-free, scalable technology framework to analyze the molecular compositions of associating complexes.

To demonstrate AMBER’s utility in characterizing higher-order chromatin associations, we analyzed nuclei isolated from the mouse cerebral cortex. At the one-dimensional level, AMBER generates reproducible chromatin fragment signals (Spearman’s correlation coefficient [SCC]: 0.95 to 0.99 among technical replicates) and agrees well with the reference ATAC-seq data^13^, recapturing major active chromatin signatures. AMBER and ATAC-seq peaks showed similar genomic distributions, with peaks unique to either method concentrated in intronic and distal regions, where shared peaks were TSS-enriched (44.4%) (**Fig. 1e** and **Extended Figs. 2d-h**). Next, we compared the high-order chromosome organization signals from AMBER with those from proximity-ligation-based Hi-C technologies. We sequenced the AMBER library derived from chromatin complexes isolated from about one million nuclei to 7.8 billion reads (PCR duplicate ratio: 78.4%); among them, 47.1% of barcodes label two or more distal chromatin fragments, and among these non-singlet barcodes, 35.5% capture multi-way accessible chromatin associating hubs (**Fig. 1f**). At the compartment level, AMBER shows good agreement with the reference Hi-C dataset of the mouse cerebral cortex^14^ (Genome-wide SCC: 0.77. **Fig. 1g, Extended Data Fig. 2i-l** and **Supplementary Table 2**). Interestingly, similar to Hi-C^14^ and Dip-C^15^, pairwise cis contacts from AMBER exhibited a steep decay in contact frequency curves, while hubs with increasing numbers of co-registered fragments showed progressively longer-range interactions (**Fig. 1h**). Therefore, AMBER captures both smaller units (e.g., chromatin fragment singlets and pairwise interactions) and larger association complexes involving multiple chromatin regions across longer genomic distances.

The ability of AMBER to resolve accessible chromatin associations across multiple distance scales prompted us to ask about the genomic features of regulatory elements engaging in interactions across chromosomes. We identified 1,409,158 beacon barcodes showing significant inter-chromosomal assignments (co-assigned locus combinations with observed over expected > 3-fold, see **Methods**) and calculated hub occupancy (HO) as the number of distinct inter-chromosomal hubs with one or more loci in non-overlapping genomic windows. Throughout the genome, 57.6% of the inter-chromosomal hubs are concentrated in a small subset of genomic domains (18,708 5-kb bins, corresponding to 3.55% of the mouse genome; **Extended Data Fig. 3a** and **Supplementary Table 3**). Interestingly, high HO regions are associated with enhancers (chromatin states: En-Sd, En-Sp, En-W, and En-Pd), particularly the Speckle-Associated Domain (SPAD)^12^, a specialized genomic region that acts as a hub for active transcription and RNA processing (**Figs. 2a, b**). Such observations cannot be explained by ATAC-seq signal density alone (**Extended Data Figs. 3b, c**). Previous observations suggested that trans-nuclear speckles preferentially localize at the nuclear center^2, 9, 15-17^; integrating AMBER with DNA-MERFISH^18^ confirmed a shift from the periphery to the center in nuclear radial positions for high HO domains (**Fig. 2c** and **Extended Data Figs. 3d-f**). Compared with peripheral counterparts, central high HO regions have higher densities of coding genes, including *Rbfox3*, which encodes an essential RNA-binding protein for nervous system development and function^19^ (**Extended Data Figs. 4a-c**). We hypothesized that if a group of genes from different chromosomes spatially organizes into regulatory hubs for co-expression, this process should benefit the assembly of protein complexes that are key to the host cell’s biological function. To test this, we performed protein-protein interaction analysis for proteins encoded in high HO region genes using the STRING database’s physical interactions^20^. The top interacting network includes proteins for neural signaling, synaptic plasticity, and synaptic transmission, which are encoded by genes located on different chromosomes, such as Chr 7 (*Grik5*), Chr 9 (*Scn4b, Grik4*), and Chr 14 (*Ptk2b, Cacna1d*) (**Fig. 2d** and **Extended Data Figs. 4d, e**). On the other hand, we did not observe genes in high ATAC-seq signal regions whose protein products are enriched for large protein-protein interaction networks (**Extended Data Fig. 4f**). Thus, AMBER captures higher-order regulatory organizations essential for cellular function.

**Fig. 2:**
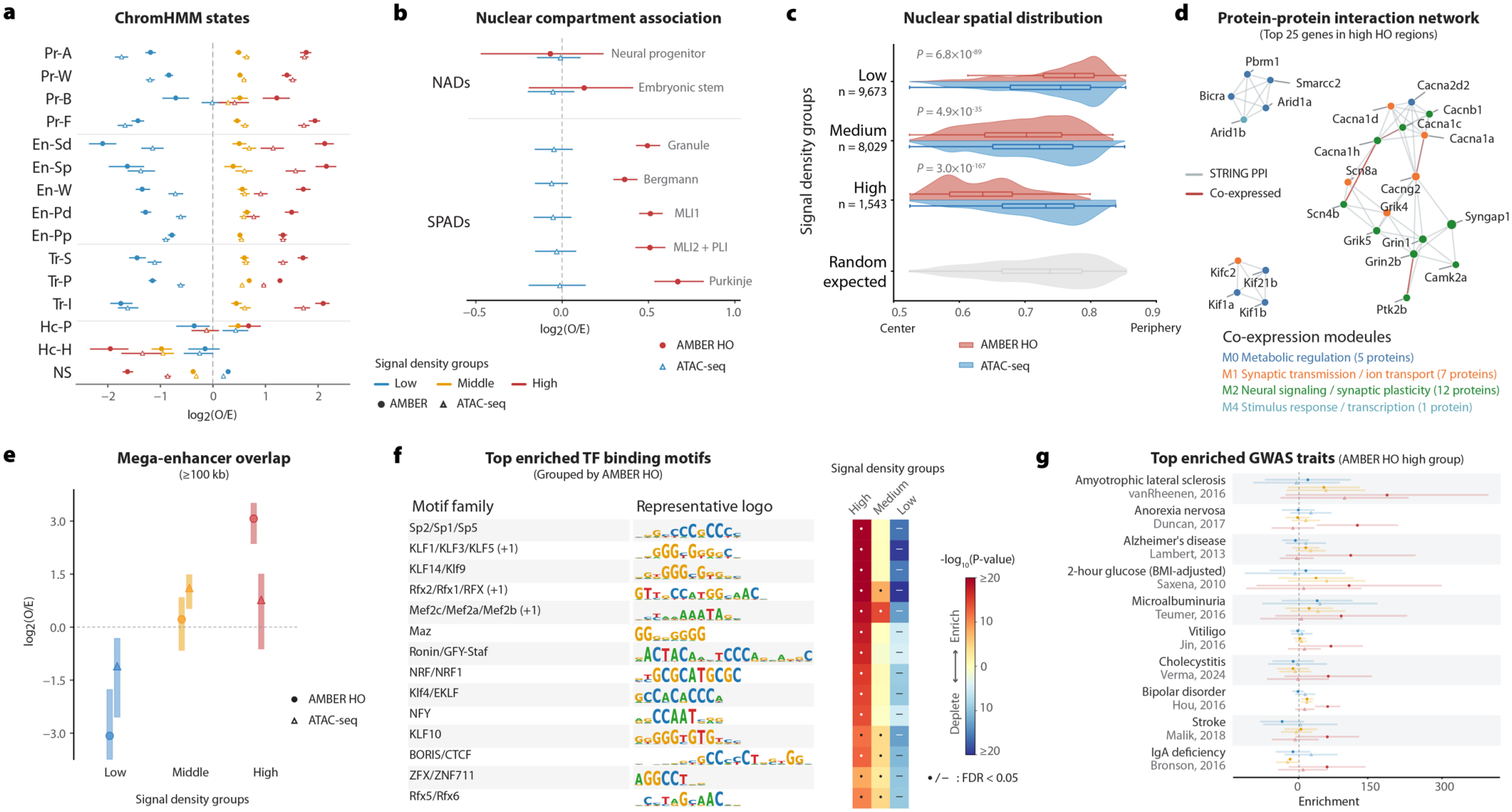
AMBER reveals inter-chromosomal regulatory hubs in the mouse cerebral cortex. **a**, Signal enrichments of ATAC-seq and AMBER in different ChromHMM chromatin state regions for high-, medium-, and low-signal density groups. Error bars show 95% CI. **b**, Enrichment of AMBER and ATAC-seq signal density by nuclear compartment association. SPAD, Speckle-associated domain; NAD, Nucleolar-associated domain. **c**, Radial position comparison of AMBER Hub Occupancy and ATAC-seq accessibility across signal density. P-value: two-sided Mann-Whitney U test. **d**, Top 25 connected genes in High HO regions from STRING protein-protein interaction network analysis. Grey edges, STRING physical interactions; red edges, co-expression from ENCODE single-nucleus RNA-seq. **e**, Enrichment of each signal density group for mega-enhancers. Bars, 95% CI. **f**, Left, HOMER known TF motifs enriched in high-density HO regions. Right, enrichment and depletion of TF motifs by signal density group. Dots mark FDR < 0.05. **g**, Enrichment of GWAS traits of human orthologues from peaks by signal density groups, ranked by Hub Occupancy.

Finally, we asked whether high HO domains are related to “Mega Enhancers” (MEs), regulatory domains showing abnormally long continuous H3K27ac signals that have previously been discovered in cerebellar granule neurons^21, 22^. Similarly, we identified MEs from mouse cerebral cortex based on a public reference H3K27ac dataset^13^. Because H3K27ac signals are strongly associated with chromatin accessibility, we first examined whether ATAC-seq signal density could strongly predict MEs: although less accessible regions are depleted of MEs, regions with the highest ATAC-seq signals do not show a stronger overlap with MEs (31% and 69%, for ATAC-seq high, and medium signal level groups, respectively; **Extended Data Fig. 5a**). In contrast, 58% of detected MEs are located in 6% of the genomic regions with high HO (**Fig. 2e, Extended Data Figs. 5a, b**, and **Supplementary Table 4**). We also found that high HO domains are strongly enriched for binding motifs of key neural transcription factors, such as RFX, MEF2, and KLF family factors (**Fig. 2f** and **Extended Data Fig. 5c**). Compared to ATAC-seq, the human genome orthologues of regulatory elements within high HO domains again display stronger associations with genetic variants of neurological disorders such as Anorexia nervosa, Alzheimer’s disease, and Bipolar disorder (**Fig. 2g** and **Extended Data Figs. 5d, e**). Together, by mapping regulatory chromatin associations, AMBER predicts disease risk loci and regulators that cannot be identified from accessibility alone.

In summary, we developed AMBER, a scalable and affordable approach to map accessible chromatin associations independent of target size and composition. By characterizing higher-order chromatin organization in mouse brain nuclei, we identified inter-chromosomal co-regulatory hubs for neural functions. Compared to proximity-ligase-based approaches^10^, AMBER resolves co-operativity and multipartite regulatory associations across a longer spatial range. The barcode expansion strategy circumvents the need for pre-manufactured barcode beads or multiple rounds of combinatorial indexing in ChIA-Drop^23^ and SPRITE^2^. While the accurate effective registration radius remains unmeasured due to technical challenges, it is a tunable parameter (e.g., by RCA duration) that enables flexible multi-scaling analysis. Here we identify co-registration by shared beacon barcodes, which support association but do not necessitate direct contact. Finally, the flexibility in barcode-seed design provides future opportunities to incorporate additional molecular modalities or biological structures such as proteins, cells, or tissue niches.

## Methods

A brief description of the AMBER experimental procedure is provided below. A more detailed, step-by-step protocol is provided on GitHub. The oligo sequences involved in the study are provided in **Supplementary Data Table 1**.

### Preparation of barcode seeds, Tn5 transposome, and chromatin complexes

#### Preparation of AMBER barcode seeds

AMBER RCA Template (0.5 µM), CircLigase ssDNA Ligase (5 U), ATP (50 µM), and MnCl_2_ (2.5 mM) were mixed in CircLigase reaction buffer (1×) and incubated for 3 hours at 60 °C, followed by 80 °C for 10 minutes, then cooled to 4 °C. The circularized fragment was verified via gel electrophoresis. Then, 10 U of Exonuclease I was added to digest uncircularized templates at 37 °C for 1 hour, followed by 80 °C for 10 mins, and cooled to 4 °C. Equimolar concentrations of circularized template and AMBER RCA Primer were combined and hybridized at room temperature for 2 hours, and adjusted to a final concentration of 0.3 µM.

#### Preparation of Tn5 transposome

Tn5 AMBER adapter A or adapter B was combined with the common pMENTs oligo (final concentration 50 µM) and annealed in a thermocycler with a program: 95 °C for 5 min, then slowly cooled down to 20 °C with a ramp of -0.1 °Cs^-1^. Then, 1 µL of this annealed adapter was mixed with 6 µL of unloaded Tn5 transposase (0.5 mg/mL), briefly vortexed, spun down, and incubated at room temperature for 30 min, followed by 10 min at 4 °C. The two types of loaded transposase were mixed in equal ratios for all tagmentation reactions.

#### Cell Preparation and crosslinking

Cultured HeLa cells or mouse cerebral cortex were prepared as a single-cell suspension and resuspended in 0.2% formaldehyde (Thermo 28906) diluted in pH 7.4 PBS for 10 mins at room temperature with end-to-end rotation mixing. The reaction was quenched with 2.5 M glycine to a final concentration of 0.2 M, incubated for 5 min at room temperature, and then placed on ice for 5 min. Cells were then centrifuged at 1,000 *g* for 10 mins at 4 °C, and the supernatant was removed. Crosslinked cells were washed once with pre-chilled 0.5% BSA in PBS solution and centrifuged at 1,000 *g* for 10 mins at 4 °C.

#### Nuclei permeabilization and *in situ* tagmentation

Cells were resuspended in 200 µL of pre-chilled OMNI Permeabilization Buffer (ATAC-RSB with 0.10% IGEPAL, 0.10% Tween20, 0.01% Digitonin, 0.5 U/µL RNaseOUT, 0.5 U/µL SUPERaseIN, and a 0.5× PI Cocktail) per 500,000 cells on ice for 5 minutes. Intact nuclei were quantified by cell counting with a small aliquot. Nuclei were centrifuged at 1,000 *g* for 10 mins at 4 °C, and the supernatant was discarded. Approximately 250,000 permeabilized nuclei were aliquoted per reaction tube and resuspended in 50 µL AMBER Tagmentation Buffer (33 mM Tris-Ac pH 7.8, 67 mM KAc, 11 mM MgAc, 16% DMF). Next, 2.3 µL of Tn5 transposome was added and gently mixed. The tagmentation reaction was carried out at 37 °C for 35 mins, with agitation at 550 rpm on a ThermoMixer (Eppendorf, EP5384000020). The reaction was terminated by adding 5 µl of 40.4 mM EDTA (Life Technologies, 15575020). Tagmented nuclei were pooled by lysis concentration, centrifuged at 1,000 g for 10 min at 4 °C, and the supernatant was removed.

#### Nuclei lysis and isolation of chromatin complexes

Tagmented nuclei were counted, and 400,000 nuclei were vigorously resuspended (∼50 pipette strokes) in 96 µL AMBER Lysis Buffer (10 mM Tris-HCl pH 8, 1 mM EDTA, 200 mM NaCl, 1% Triton X-100, 0.75% SDS, 0.75% SDOC, 1× EDTA-free Protease Inhibitor Cocktail) to release chromatin complexes. Lysis was carried out at 37 °C for 1.5 h with shaking at 800 rpm (ThermoMixer, Eppendorf). Cell lysis was validated by cell counting of a small aliquot under the microscope. Next, 64 µL of No-salt AMBER Washing Buffer (10 mM Tris-HCl pH 7.5, 1 mM EDTA, 0.1% Tween-20) and 20 µL of 10% IGEPAL CA-630 (Sigma, I8896) were added to each lysate tube and mixed. Ten microliters of Streptavidin C1 beads were washed 3 times with Low-salt AMBER Washing Buffer (10 mM Tris-HCl pH 7.5, 1 mM EDTA, 0.1% Tween-20, 150 mM NaCl), then added to the tubes and incubated for 2 hrs at room temperature with end-to-end rotation. The tube was gently spun down, placed on a magnetic separation rack (New England Biolabs S1515S), and the supernatant was carefully discarded. Beads were washed twice with 200 µL of Low-Salt AMBER Washing Buffer. Beads were resuspended in 100 µL of 1× KAPA HiFi HotStart Uracil+ ReadyMix Kit (Roche, KK2802) at 72°C for 10 minutes at 1,050 rpm on a ThermoMixer for gap-filling and heat-displacement of Tn5-derived ME oligo. Beads were then washed 1× with Low-salt AMBER Washing Buffer, followed by 3 washes with No-salt AMBER Washing Buffer. The washed beads were resuspended in 10 µL of nuclease-free water. Elution was conducted by slowly heating to 80 °C (with a ramp rate of 0.5 °Cs^-1^) and holding for 45 seconds. Beads were placed on a magnet and supernatant containing chromatin complexes was transferred to a new tube. Concentration was determined using a spectrophotometer. A 1 µL aliquot was set aside for decrosslinking. Qubit quantification and PCR amplification were used as quality-control proxies.

### AMBER procedures

Hydrogel formation. The AMBER hydrogel master mix was prepared on ice to prevent gel polymerization. For a 10 µL reaction, the following reagents were combined to their final concentrations: chromatin fragments (∼400 ng), AMBER barcode seeds (17.4 nM), 4-arm PEG acrylate 10K MW and HS-PEG-SH 3.4K MW (combined 9.5% w/v), Phi29 DNA polymerase (0.83 U), Phi29 reaction buffer (1×), DMSO (4.2%), BSA (4.2 mg/mL). The gel was left to incubate at 30 °C for 30 minutes with a heated lid (100 °C).

#### Seed expansion and chromatin indexing

dNTP mix (2.25 µL of 5.5 mM dNTP, 4.4 × CutSmart Buffer) was gently overlaid on top of the polymerized gel and left to submerge for 2 hrs at 4 °C, then high-efficiency rolling circle amplification was conducted at 37 °C for 2 hrs to generate multiple copies of the seed oligo in-gel, and Phi29 was denatured at 65 °C for 20 mins. A dual-enzyme cutting mix (4.5 µL of 15 U HaeIII and 15 U Nb.BsrDI in 3.3 × CutSmart Buffer) was then overlaid, and the cutting reaction was carried out at 37 °C for 10 hrs; the enzymes were then denatured at 80 °C for 20 mins. HaeIII cleaves single-molecule spatial barcodes containing the same beacon barcodes into multiple ssDNA, whilst Nb.BsrDi and heating expose the cognate hybridization region on the genomic DNA. Ligation mix (5.5 µL of 4,500 U T7 DNA Ligase in 1.45× StickTogether DNA Ligase Buffer) was then overlaid and incubated at 20 °C for 20 hours to facilitate ligation between gDNA adapter sites with cleaved ssDNA beacon barcodes. Following incubation, the gel was placed on ice, the supernatant was removed, and it was washed once with chilled PBS to remove residual reagents.

#### Gel denaturation

To denature the hydrogel, 40% of the gel volume in pre-chilled AMBER Gel Denaturation Buffer (457.5 mM KOH, 100 mM EDTA, 42.5 mM DTT) was added and mixed to degrade gel crosslinks via alkaline hydrolysis. Thorough denaturation was ensured by incubation for 2 hrs at 4 °C, with intermittent pipetting twice. The solution was then neutralized with 12 µL of AMBER Gel Neutralization Buffer (600 mM Tris-HCl pH 7.5, 305 mM HCl) at room temperature and spun down.

#### DNA extraction, library amplification, and sequencing

Ten microliters of pre-washed Streptavidin C1 beads and 80 µL Low-salt AMBER Washing Buffer were added to the dissolved gel and incubated with end-to-end rotation for 1 hr at room temperature to recollect complexes. After binding and washing with Low-salt AMBER Washing Buffer for 3 times, the dried beads were resuspended in 20 µL of PBS with 1.2 U of Thermolabile Proteinase K (NEB, P8111) and de-crosslinked at 37 °C, 1,400 rpm on a ThermoMixer for at least 8 hrs. The Thermolabile Proteinase K was deactivated by heating at 55 °C for 15 mins at 1,000 rpm. The beads were isolated via magnet rack isolation and supernatant used directly for PCR (72 °C for 5 min; 98 °C for 30 s, 98 °C for 20 s, 63 °C for 20 s, 72 °C for 1 min; 72 °C for 2.5 min; hold at 12 °C) with separate Illumina-indexed forward and reverse primers (Nextera N5 and N7) for each AMBER gel, in 1× KAPA HiFi HotStart Uracil+ ReadyMix Kit. The PCR included a 5 mins 72 °C pre-extension step to prepare the opposing strand of the ssDNA beacon barcode monomer for PCR. An initial 9 cycles were conducted, and a 2 µL aliquot underwent qPCR to determine the required cycles to reach 20 nM. The final library was cleaned using 0.8 × SPRI beads and eluted in 15 µL nuclease-free water. Sequencing was conducted on the NovaSeq X platform (Illumina) with paired-end read 150 + 150 cycles, and on the Ultima Genomics UG100/UG200 sequencer.

### Data analysis

Barcode hub reconstruction. Unaligned reads were filtered for ligation bridge (GCATGCGG) with one mismatch allowed, which verifies ligation of the ssDNA beacon barcode to chromatin in gel. The randomized beacon barcode was extracted, and barcodes that violated the degenerate design (BNNBNNDNNBNNBNN) were removed. Adapter sequences, barcode, and clamp were trimmed with cutadapt (v4.4; error rate 0.18, quality cutoff Q15, minimum length 20 nt), and the remaining Tn5-targeted accessible DNA was aligned to mm10 with bwa-mem2 (v2.2.1). Alignments were restricted to primary alignments with MAPQ ≥20, and those overlapping RepeatMasker elements (UCSC, milliDiv <140) and the ENCODE mm10 blacklist (v2) were removed. Beacon barcodes were used to group genomic hubs with UMI-tools (directional method, Hamming distance threshold 2). Within each hub, fragments starting within 147 bp (nucleosome distance) of a retained fragment on the same chromosome were collapsed as duplicates.

#### Hub interaction scaling correction

Because the number of pairwise contacts in a hub grows quadratically with its number of fragments, large hubs would disproportionately dominate contact frequency maps. To correct this, each hub of n fragments (n ≥ 2) was decomposed into its n(n-1)/2 pairwise contacts, and each contact was assigned a weight of 2/n, so that a hub contributes a total weight of n−1 and scales linearly with its size. Contacts were restricted to chromosomes 1-19 and X. Weighted pairs were binned at 25 kb with cooler (cload pairs), aggregated to resolutions from 100 kb to 5 Mb with cooler zoomify, and normalized by iterative correction (ICE) using cis contacts only.

#### Statistically significant multi-way interaction hub permutation filtering

To identify significant multi-way hubs that recur more often than expected by chance, each hub was reduced to a signature: the set of unique 1-Mb genomic bins containing its fragments. Hubs spanning fewer than two bins were excluded, and hubs with identical signatures were counted to give an observed frequency. Signatures observed at least three times were tested against a permuted null (n = 10,000) restricted to a “universe background” of 1 Mb bins covered by bulk cortex ATAC-seq or AMBER fragments, so that permuted hubs could only fall in regions the assay can detect. For each permutation, a cis signature was randomly relocated within its chromosome with its internal spacing preserved, and each bin of a trans signature was resampled from the universe bins on the same chromosome. The null frequency of a permuted signature was its observed frequency in the dataset. Expected frequency was the mean across permutations, enrichment (O/E) was observed over expected, and the empirical P value was the fraction of permutations with a frequency at least as high as observed. Signatures with O/E >3 (or zero expected frequency) and P < 0.05 were retained as significant hubs for analysis.

#### Seed Reporter collision analysis

Seed and Reporter collisions produce double-stranded, sequenceable products by proximity hybridization, with a structure as described: Forward PCR handle-Seed barcode-Flank-Complementary hybridization region-Flank-Reporter barcode-Reverse PCR handle. After sequencing, reads with a mean Phred quality score of ≥20 and matching the barcode design were retained and deduplicated. Each unique pair was counted as one collision event. Seed barcodes supported by ≥2 reads were taken as confirmed, and the collision rate was calculated as the fraction of confirmed Seed barcodes paired with two or more distinct Reporter barcodes.

#### Peak calling and annotation

Peaks were called using MACS3 with the following parameters: (--nomodel --shift - 75 --extsize 150 -q 0.01 --nolambda). Genomic feature annotation and distribution were performed with ChIPseeker against GENCODE vM25 TxDb, and promoters were defined as ±3 kb relative to the transcription start site.

#### Stratum-adjusted correlation coefficient

AMBER replicates were preprocessed by quantile transformation, and stratum-adjusted correlation coefficients were determined by HiCRep. This method transforms the features to follow a normal distribution, enabling comparisons between independent features. Because AMBER was designed to resolve open regulatory region interactions, direct correlation scores with genome-wide interaction mapping methods such as Hi-C and SPRITE are inappropriate here. Reproducibility between AMBER gel libraries was assessed with HiCRep stratum-adjusted correlation coefficients (SCC). Sub-libraries were first thinned to equal depth by randomly retaining whole hubs, and cis contacts were taken as all fragment pairs within a hub with weight 2/n. This method smooths each contact map and computes Pearson correlations within strata of genomic distance, then combines them with stratum-specific weights into the SCC, which controls for the distance dependence of contact frequency.

#### Saddle plots and Eigenvector calculations

Compartment scores were computed with cooltools eigs_cis on cis-only ICE-balanced maps, per chromosome (1-19 and X), taking the first eigenvector (E1) with the sign set by GC content of mm10 (positive = A, negative = B). O/E maps used the per-diagonal mean of balanced contacts as the expectation (expected_cis, unsmoothed), log2-transformed with a floor of 0.125, with masked Gaussian smoothing (sigma 1 bin for AMBER, 0.5 for Hi-C). For saddle plot analysis, bins were ordered by Hi-C E1 eigenvector, and cis and trans O/E were averaged per group pair with cooltools saddle. Saddle strength score was calculated as (AA + BB) / (AB + BA), computed from corner groups covering the top and bottom 20% of the E1 range.

#### Contact frequency decay curves

Contact probability decay curves were generated with cooltools expected_cis at 10-kb resolution on cis-only balanced maps. Curves were smoothed on log-log axes and divided by their first valid genomic separation point (20 kb). Comparative method datasets were retrieved from Hi-C (4DNFI2H23JBK) and Dip-C (GSE162511).

#### Hub Occupancy calculation

Hub occupancy (HO) of a genomic region was defined as the number of significant inter-chromosomal hubs with at least one fragment in the bin. Fragments per hub were counted at most once per bin. Bins with HO >0 were ranked and divided into Low, Middle, and High groups at the median and at the knee of the ranked HO curve above the median (Kneedle; HO capped at the 99.9th percentile). ATAC-seq groups of the same sizes were formed by ranking bins by cortex ATAC-seq signal.

#### Chromatin state annotation

Chromatin states were annotated with the published ENCODE3 15-state ChromHMM segmentation of mouse postnatal day 0 forebrain (mm10; see Data Availability). For each 100-kb bin, the base pairs covered by each state were summed. Group enrichment was observed state coverage divided by the coverage expected from chromosome-matched random bins (all QC-passing 100-kb bins, including HO = 0), shown as log2(O/E) with 95% intervals from 499 chromosome-stratified 1-Mb block bootstrap replicates.

#### Nuclear compartment association calculation

NAD and SPAD region (see Data Availability) coordinates were merged and intersected with 100-kb bins. Observed overlap per group was divided by the expectation from bins matched by chromosome, GC-content quintile, and ATAC-seq quintile, shown as Log_2_(O/E) with 95% intervals from 499 chromosome-stratified 1-Mb blocks bootstrap replicates.

#### Genomic analysis of nuclear radial positions

We measured the radial position of each DNA-MERFISH spot by normalizing its distance from the nuclear center (derived from the mean of 988 genome-wide spots); cells with a minimum of 600/1,976 spots were retained for this analysis to the surface convex hull (defined by all detected spots) (**Extended Data Fig. 3**). To annotate genomic regions from AMBER, we estimated radial scores by linear interpolation between two flanking MERFISH probes, and assigned a confidence score inverse to the genomic distance from the nearest imaged locus. Regions of confidence ≥0.4 (flanking probes ≤3 Mb apart) were retained for radial analyses.

#### Protein-protein interaction network analysis

Protein-coding genes overlapping the 100 kb bins of each signal density group formed the gene set. Co-expression was computed from single-nucleus RNA-seq (See Data Availability) and partitioned into modules by Leiden clustering. Genes in a set with a co-expression partner in the same set were linked by STRING v12.5 mouse physical interactions (combined score ≥700).

#### Mega Enhancers assignment and radial annotation

Mega enhancers were defined following a strategy described previously. H3K27ac ChIP-seq peaks were stitched together (<12.5 kb gap distance), and regions exceeding 50 kb, 100 kb, and 200 kb cutoffs were classified as MEs. Protein-coding genes overlapping mega-enhancer bins were assigned to a subpopulation based on their bin’s radial position.

#### HOMER and GO enrichment

Motif enrichment was assessed with HOMER. 200bp windows centered on ATAC-seq peak summits were assigned to the signal density group of their 100-kb bin and scanned for known motifs (homer2 find, original thresholds, both strands). The windows of each group were tested against all remaining windows by a two-sided Cochran-Mantel-Haenszel test stratified by chromosome, GC content and CpG count, with Benjamini-Hochberg adjustment.

#### GWAS trait enrichment analysis

We compared signal density groups with genome-wide association studies (GWASs) of human phenotypes, as in previous study^24^. Briefly, mouse cortex ATAC-seq peaks were intersected with the 100-kb bins of each group (Extended Data Figure 5d), and the intersected intervals were converted from mm10 to hg38 with liftOver (UCSC mm10ToHg38 chain; ‘-minMatch=0.5’), retaining uniquely mapped autosomal intervals with at least 50% of bases remapped. Estimation of enrichment coefficients of each trait compared to the background control was performed using the LDSC software (https://github.com/bulik/ldsc).

## Data availability

Raw sequencing data generated in this study have been deposited in NCBI GEO and will be available after revision. External datasets were downloaded from GEO with the following accession numbers: mouse brain Dip-C (GSE162511), cerebellar granule neuron SPRITE (GSE339045), and nucleolus-associated domains in mouse embryonic stem cells and neural progenitor cells (GSE150822). Speckle-associated domains (SF3A66 peak calls in adult mouse cerebellar cell types; Takei et al., 2025) were obtained from the CaiGroup/dna-seqfish-plus-multi-omics GitHub repository. ENCODE (https://www.encodeproject.org/) with the following accession numbers: mouse frontal cortex ATAC-seq (ENCSR889WQX), mouse cerebral cortex histone ChIP-seq for H3K27ac (ENCSR491RBV), H3K4me3 (ENCSR943LBN) and H3K4me1 (ENCSR000CAI); mouse cerebral cortex TF ChIP-seq for CTCF (ENCSR644VYX); mouse postnatal day 0 forebrain histone ChIP-seq for H3K27me3 (ENCSR070MOK), H3K9me3 (ENCSR093DWU), H3K4me2 (ENCSR906UJW), and H3K9ac (ENCSR369RBO) and mouse cortex single-nucleus RNA-seq (ENCSR442EUE). The mouse postnatal day 0 forebrain 15-state ChromHMM annotation (ENCODE3) was obtained from the UCSC Genome Browser (mm10, encode3RenChromHmmForebrainP0). Protein-protein interactions were obtained from STRING v12.5 (mouse, physical links; https://string-db.org/). 4DN Data Portal (https://data.4dnucleome.org/) with the following accession numbers: mouse cerebral cortex Hi-C (4DNFI2H23JBK) and mouse cortex DNA-MERFISH (4DNESPE924IP).

## Code availability

The code used in this study is available on GitHub (https://github.com/czhulab/AMBER).

## Acknowledgements

We thank QB3 MacroLab for the Tn5 enzymes. We thank Jessica Young for assistance with illustrations. C.Z. is supported by Weill Cornell Medicine and New York Genome Center startup funds, National Institutes of Health (NIH)/National Institute of General Medical Sciences (grant no. DP2GM154011), NIH/National Cancer Institute (grant no. R61CA309705), and NIH/National Human Genome Research Institute (grant nos. R00HG011483, RM1HG011014), and the MacMillan Center for the Study of the Noncoding Cancer Genome at the New York Genome Center.

## Author contributions

R.T. and C.Z. conceived the study and analyzed the data. R.T. developed the AMBER protocol and generated the data. R.T. and C.Z. wrote the manuscript with input from J.S. C.Z. supervised the study.

## Ethics declarations

The authors declared no conflicts of interest.

---

**Extended Data Fig. 1:**
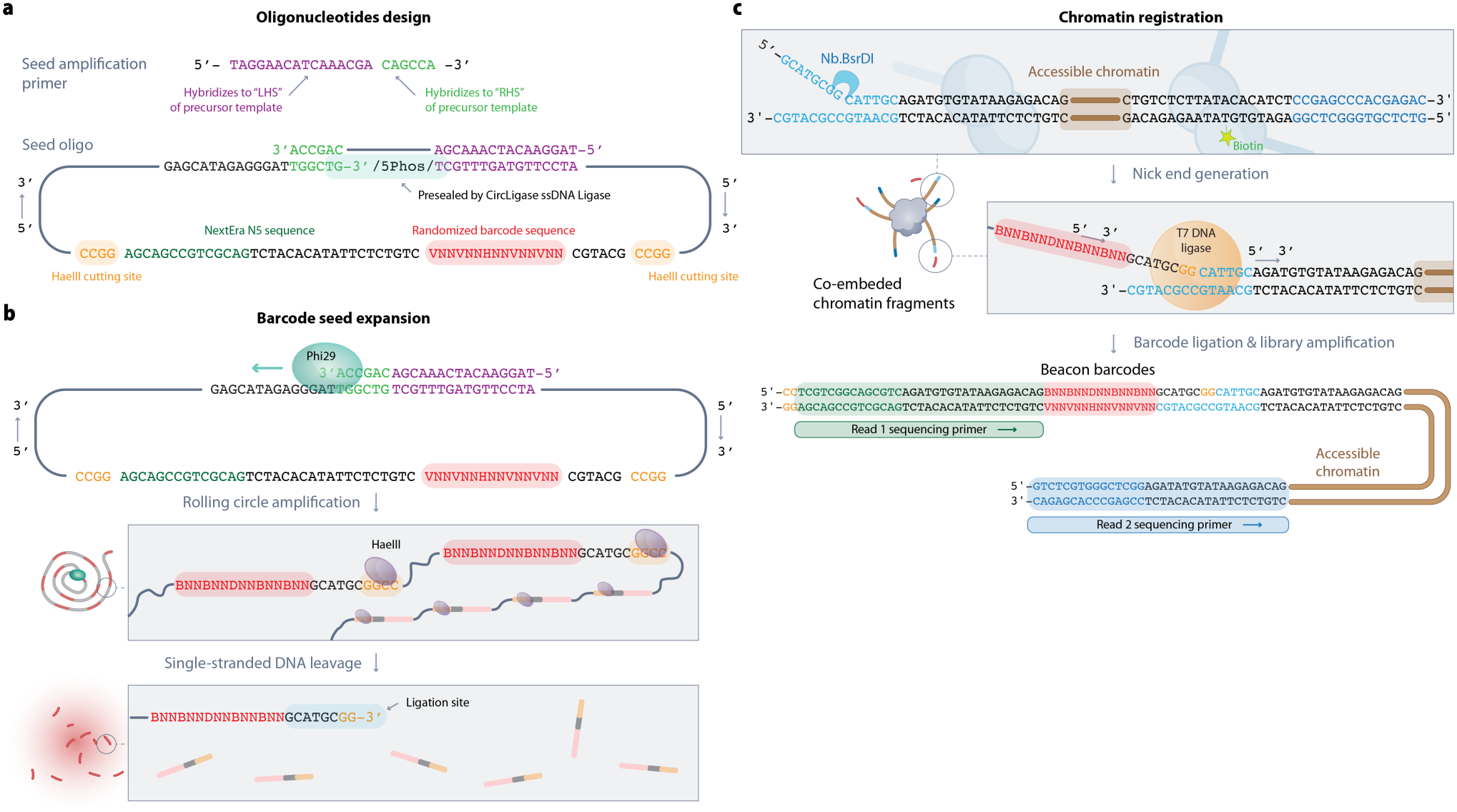
Experimental design and workflow of AMBER. **a**, Design and preparation of seed oligos. Seed amplification primer (top) is hybridized to a pre-circularized Seed Oligo (bottom) containing a randomized barcode sequence (red). **b**, Beacon barcode seed expansion and release. Top: Phi29 polymerase generates multiple copies of the seed oligo in a single strand, including randomized barcodes. Bottom: HaeIII cleaves the repeated motif to generate single-stranded monomer barcodes. **c**, Chromatin registration. Nb.BsrDI nicks the Tn5-placed adapter on accessible genomic chromatin. T7 DNA Ligase co-registers nearby monomer barcodes.

**Extended Data Fig. 2:**
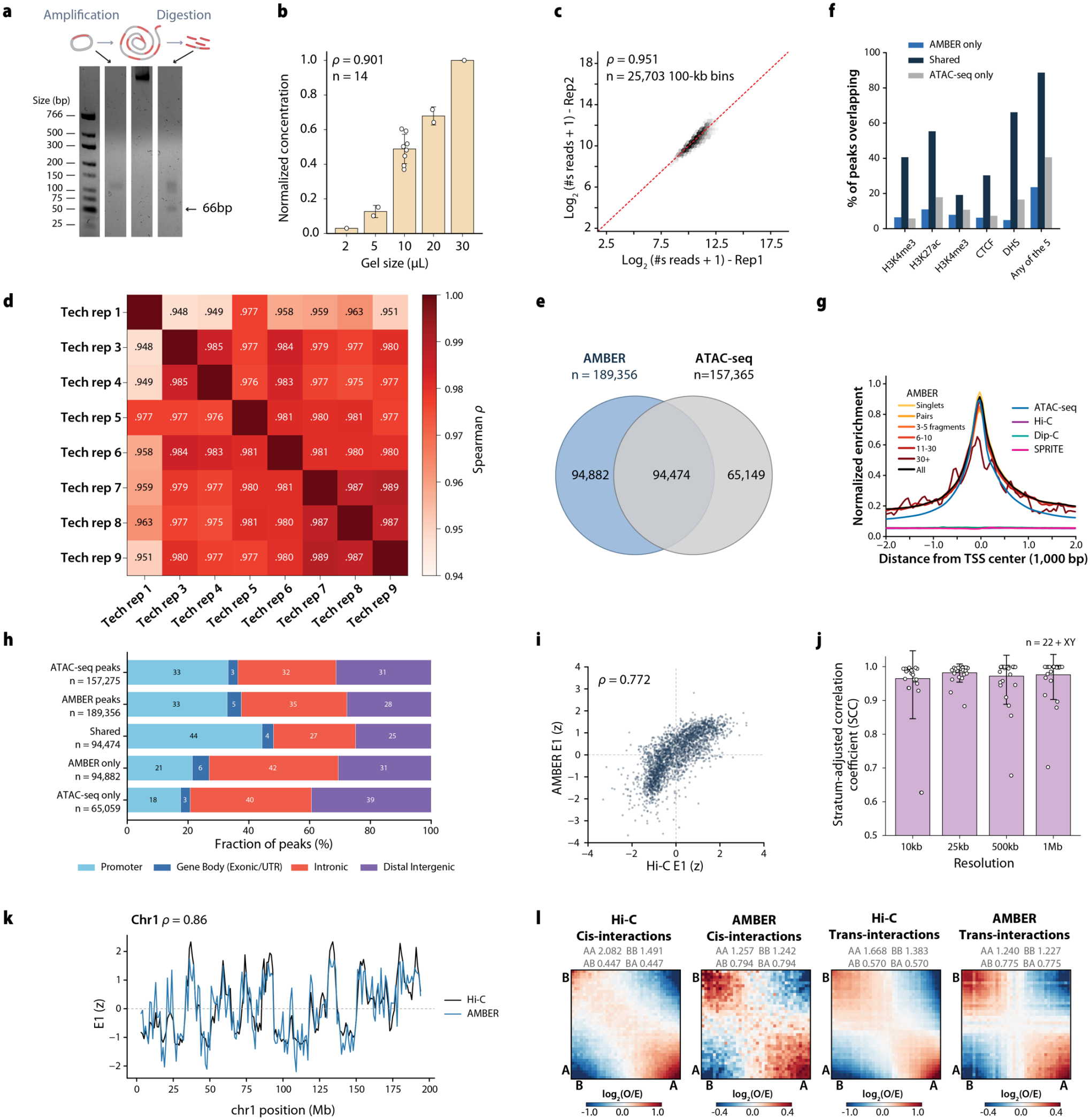
Validation of AMBER. **a**, Gel electrophoresis showing amplification and digestion of AMBER beacon barcodes. From the second lane: Left, Circularized seed oligo with random template and primer. Middle, Beacon barcode after seed expansion. Right, digested products of varying tandem repeats. **b**, Evaluation of AMBER hydrogel-reaction scalability by post-AMBER library amplification (normalized concentrations) versus input hydrogel sizes at equal concentrations. **c**, Correlation scatterplot of read counts (Log2[fragments+1]) between two technical replicates. **d**, Heatmap of showing the correlation among AMBER technical replicates. **e**, Venn diagram overlap of AMBER (blue) and ATAC-seq (grey) peaks. **f**, Percentage of peaks, as stratified in e, that overlap regulatory features H3K4me3, H3K27ac, H3K4me1, CTCF, DHS, or any of the five. **g**, Comparative enrichment plot of sequence depth-normalized reads for AMBER and hub sizes (red gradient) alongside ATAC-seq (blue), Hi-C, Dip-C, SPRITE, plotted as signal coverage surrounding ±2 kb transcription start sites. **h**, Genomic feature distributions of open chromatin peaks. **i**, Scatter plot of genome-wide correlation of A/B compartments at 1 Mb resolution. **j**, Technical replicate Stratum-Correlated Coefficient (SCC) scores at varying contact frequency resolutions. SCC bars represent mean SCC scores of all chromosomes with S.E.M. **k**, Compartment eigenvector tracks (1 Mb) as displayed in Fig. 1g, along chromosome 1, displayed as 1D overlaps. **l**, Compartment saddle plots for Hi-C (left) and AMBER (right), in cis (top row) and trans (bottom row).

**Extended Data Fig. 3:**
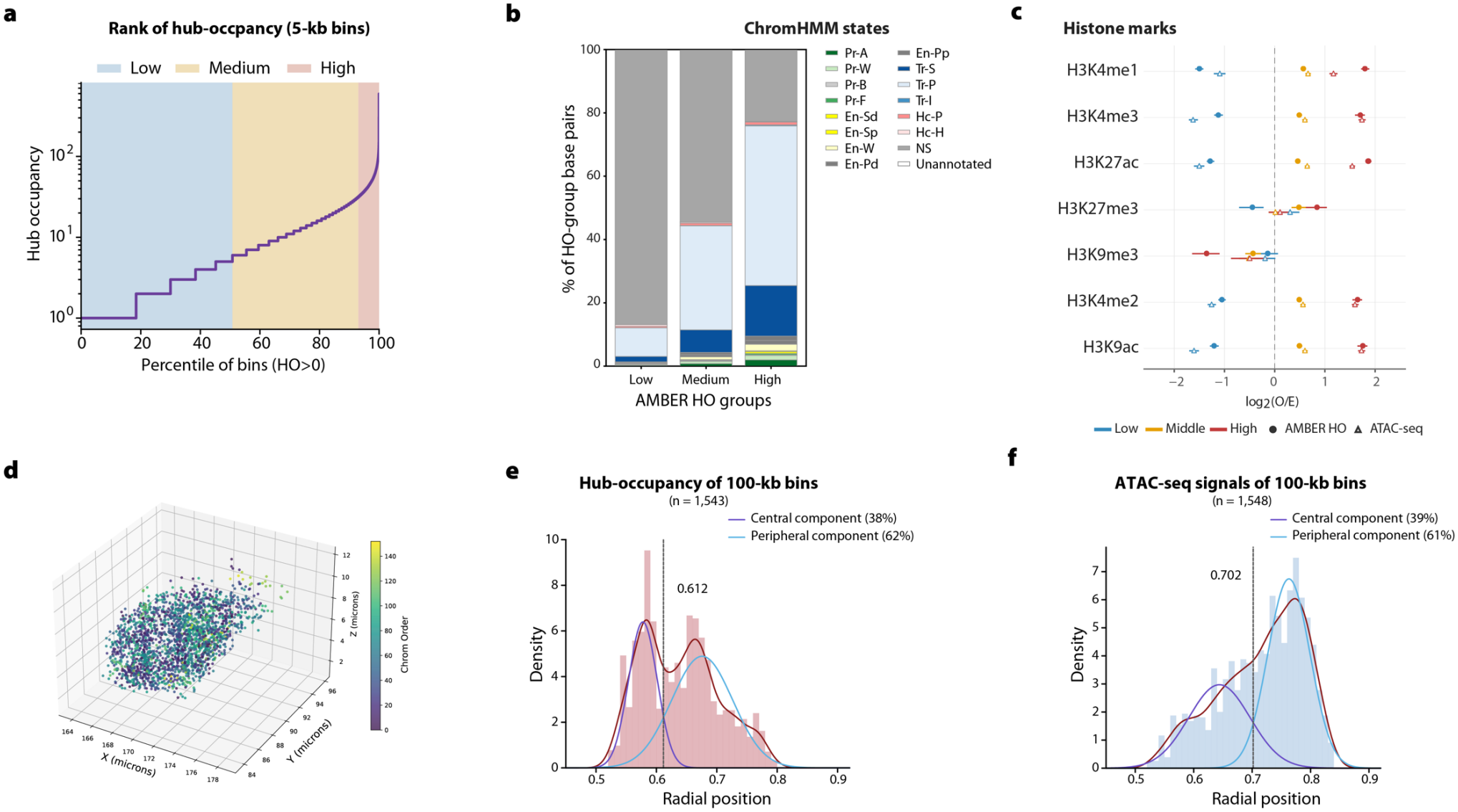
Distributions of AMBER inter-chromosomal interaction hubs. **a**, Distribution of Hub Occupancy signal coverage across the genome in 5 kb bins. **b**, ChromHMM state annotation for Hub Occupancy grouping. **c**, Comparison of overlap groupings between AMBER Hub Occupancy and ATAC signal strength on ChIP-seq histone marks. **d**, 3-D reconstruction and demonstration of DNA MERFISH chromatin localization spots. **e** and **f**, Gaussian mixture modeling of radial positions as inferred by MERFISH of High HO (e) and high ATAC-seq strength (f) regions.

**Extended Data Fig. 4:**
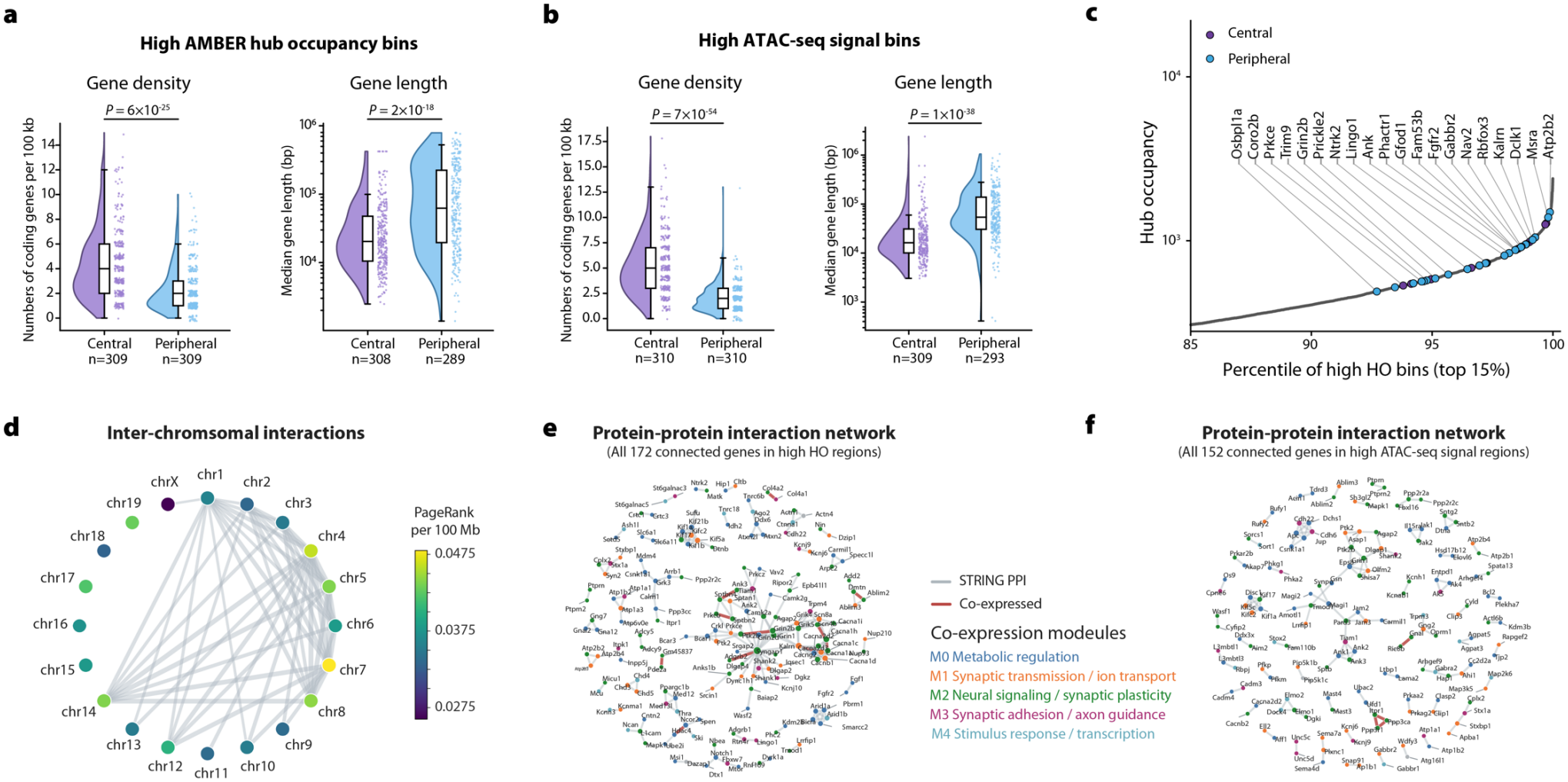
Characterization of genes associated with inter-chromosomal hubs. **a** and **b**, Violin dot plots comparing gene density and gene length between the most centrally and peripherally localized (bottom and top 20% radial positions, respectively) for high Hub Occupancy (a) and ATAC-seq signal density (b) chromatin regions. P-value: two-sided Mann-Whitney U test. **c**, Ranked zoom-in at top 15% high-density Hub Occupancy regions. Bubble points mark the highest occupancy bin of each labeled gene overlapping a mega enhancer region (> 100 kb). Colors mark radial subpopulation leaning (central, purple; peripheral, blue). **d**, Network of significant inter-chromosomal interactions, linked by co-occurrence. **e** and **f**, Protein-protein interaction networks of the most connected genes derived from high Hub Occupancy (e) and ATAC-seq signal density (f) chromatin regions. Grey edges: STRING physical interactions; Red edges: co-expression from public single-nucleus RNA-seq data.

**Extended Data Fig. 5:**
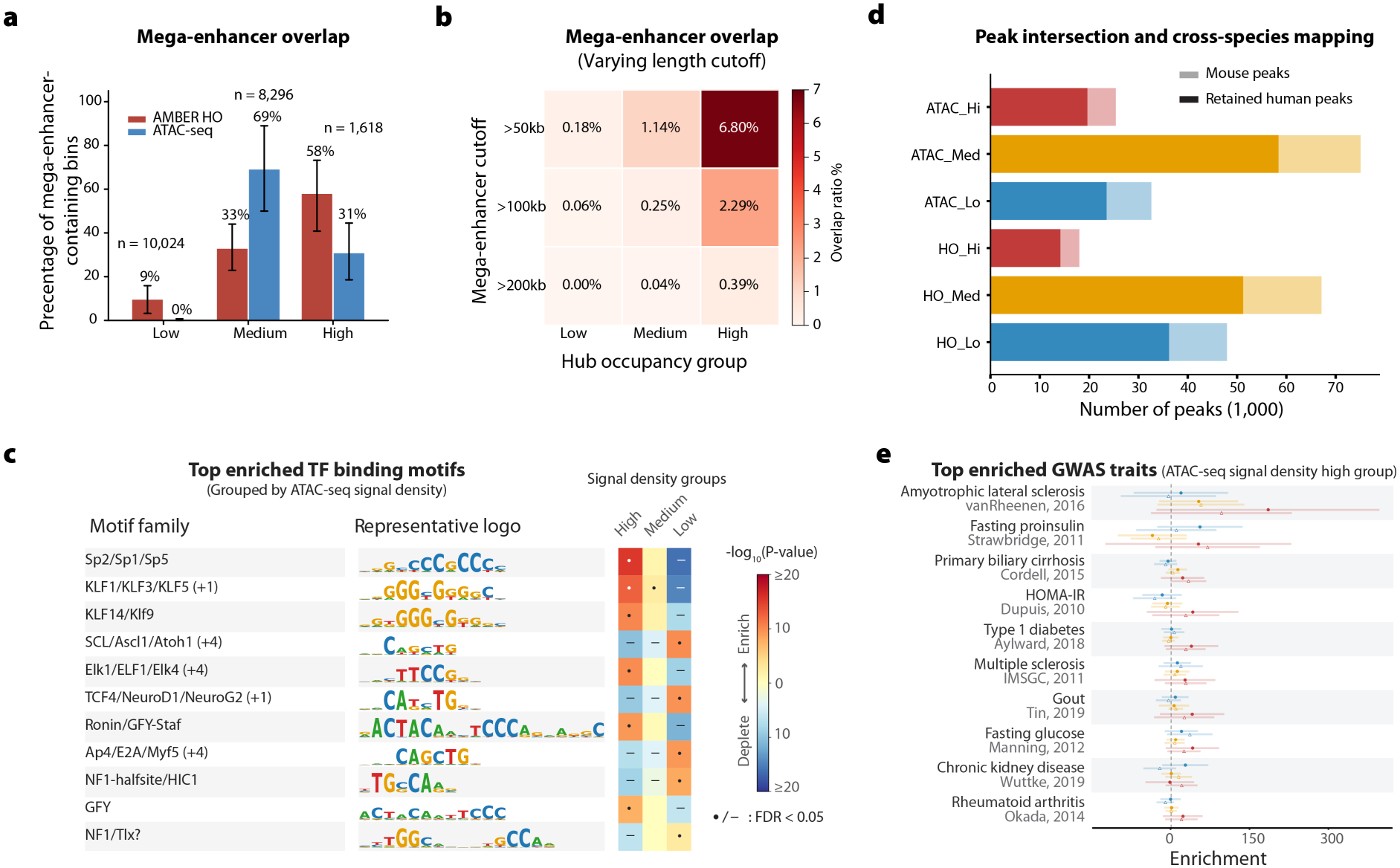
AMBER detects regulatory hubs associated with mega-enhancers. **a**, Percentage of regions in each signal density group containing mega-enhancer overlap, for Hub Occupancy (red) and ATAC-seq (blue) groups. Error bars: 95% CI. **b**, Heatmap showing the overlap between hub occupancy density regions and mega-enhancer regions at three size cutoffs. **c**, Left, HOMER known TF motifs enriched in high-density ATAC-seq regions. Right, enrichment and depletion of TF motifs by signal density group. Dots mark FDR < 0.05. **d**, Intersection of mouse peaks (light) retained after liftover to hg38 (dark) for each signal density group. **e**, Enrichment of GWAS traits of human orthologues from peaks by signal density groups, ranked by ATAC-seq high signal density regions.

## Notes

### Competing Interest Statement

The authors have declared no competing interest.

